# Microtubules in the coenocyte *Phytophthora* function in nuclear positioning and sustaining tip growth

**DOI:** 10.1101/2025.10.02.679959

**Authors:** Michiel Kasteel, Kiki Kots, Johan van den Hoogen, Edouard Evangelisti, Francine Govers, Tijs Ketelaar

## Abstract

The microtubule cytoskeleton consists of dynamic intracellular filaments and is involved in numerous processes, ranging from nuclear division to intracellular transport. Some of these microtubule-mediated processes are conserved in all eukaryotic lineages while others are specific for certain groups of organisms. Here, we focus on the microtubule cytoskeleton of oomycetes in the genus *Phytophthora*, a group of harmful plant pathogens. For visualizing microtubule organization and dynamics, we generated transgenic *Phytophthora palmivora* lines expressing GFP-tagged α-tubulin. Besides a conserved localization in the mitotic spindle, we observed cytoplasmic microtubules originating from microtubule-organizing centers associated with nuclei in the coenocytic hyphae. These dynamic microtubules initiated long-lasting, antiparallel connections with microtubules originating from adjacent nuclei. After mitosis, these microtubules rapidly increased in length while maintaining their antiparallel interaction. Frequent buckling events suggest that microtubule-based force generation plays a role in nuclear spacing. This idea was strengthened by erratic nuclear motility and positioning in hyphae exposed to the microtubule depolymerizing drug oryzalin. Besides aberrant nuclear positioning we also observed defects in hyphal growth; in oryzalin-treated hyphae lacking microtubules, tip growth was less sustained than in non-treated hyphae. This suggests that microtubules radiating into the hyphal tip from the apical nucleus have a function in sustaining tip growth. Altogether, this study provides novel insights in the localization, dynamics and functions of the microtubule cytoskeleton in the coenocytic *Phytophthora* hyphae.

## Introduction

Microtubules are filamentous protein polymers that are key components of the eukaryote cytoskeleton (Wickstead & Gull 2011). They are essential for various cellular processes, including vesicle trafficking, mitosis and cilia- or flagellar-based motility (Erickson 2007, Xiang & Plamann 2003). Microtubules are hollow, 25 nm wide tubes assembled from heterodimeric subunits consisting of α- and β-tubulin (Desai & Mitchison 1997). The uniform orientation of these heterodimers within the microtubule confers intrinsic polarity, resulting in distinct plus (+) and minus (-) ends. Microtubules alternate phases of polymerisation and depolymerisation, a phenomenon referred to as dynamic instability. This process is primarily observed at the + end. The - end is stabilized by, e.g., its attachment to a microtubule organizing center (MTOC), or shrinks under physiological conditions (Horio & Murata 2014). Due to their dynamic instability and their association with proteins that can modulate microtubule dynamics, so-called Microtubule-Associated Proteins (MAPs), the microtubule cytoskeleton can rapidly restructure in response to intra- and extracellular cues. MAPs include proteins involved in nucleating, stabilizing and bundling microtubules. Motor proteins, dyneins and kinesins (Wade 2009), can have a role in directionally translocating cargo along microtubules, but can also function in structuring the microtubule cytoskeleton by, e.g., sliding microtubules apart or by inhibiting microtubule polymerisation (Bodakuntla et al 2019). Although some microtubule-mediated processes are conserved, others differ between evolutionary groups (Gardiner 2013). Functional differences may be accompanied by distinct microtubule organizations: In plant cells, for instance, microtubules form an acentrosomal cortical array that guides cellulose-depositing enzyme complexes (Murata et al 2005) and assemble into a cortical band during the mitotic preprophase that templates the division plane (Mineyuki 1999). Many fungi possess MTOCs (referred to as spindle pole bodies) which, unlike animal MTOCs (called centrosomes), lack centrioles (Jaspersen 2021).

Our research focuses on oomycetes, filamentous microbes which can be saprophytic but are primarily known as devastating pathogens of plants, animals, insects and other microbes (Govers, 2024; Thines, 2018). They exhibit a morphology similar to fungi, characterized by filamentous hyphae with tip growth and spores for dispersal. Nonetheless, oomycetes are classified within the Stramenopiles lineage along with brown algae and diatoms (Levesque 2011), and are thus phylogenetically distant from Unikonts, the supergroup that harbours the fungi (Koonin 2010). Among the oomycetes, the genus *Phytophthora* comprises many devastating plant pathogens of agriculturally and ecologically important crops and plants. Examples include *Phytophthora infestans*, the causal agent of late blight in potato and tomato (Fry 2008), and *Phytophthora palmivora*, a broad-host range pathogen on, e.g., cacao and oil palm (Drenth & Guest 2013). Our research aims at unravelling cellular processes in *Phytophthora* and exploiting the acquired insights to identify novel potential targets for disease control. In this study we focus on the microtubule cytoskeleton of these devastating pathogens.

Studies on the microtubule cytoskeleton in oomycetes are scarce, limited to a few species and solely based on localization studies in fixed material. Immunolocalization showed that microtubules in *Saprolegnia ferax* are mostly oriented parallel to the long axis of the hyphae (Heath & Kaminskyj 1989) while electron microscopy studies in the same species revealed that microtubules originate from MTOCs associated with the nuclear membrane (Heath & Greenwood 1968) and are most abundant in the nuclei-rich zones. Transmission Electron Microscopy revealed that in *P. infestans*, cytoplasmic microtubules appear in bundles of approximately ten (Temperli et al 1990). In *S. ferax*, it was found that microtubules rarely extend into the apex of hyphae (Heath & Kaminskyj 1989). Additionally, exposure to microtubule depolymerizing drugs resulted in slower hyphal growth, reduced straightness and more frequent branching (Heath et al 2000). Since microtubule depolymerization did not fully inhibit growth, it is unlikely that the microtubule cytoskeleton in oomycete hyphae mediates the transport of vesicles containing cell wall precursors, with that role likely being fulfilled by actin (Ketelaar et al 2012). However, since the directionality of hyphal growth becomes more erratic upon microtubule depolymerization, it was suggested that microtubules have a role in maintaining growth directionality (Heath et al 2000). A similar role for microtubules has been implicated in tip-growing cells of other organisms, including fungal hyphae (Riquelme et al 1998), root hairs in *Arabidopsis thaliana* (Ketelaar et al., 2002) and moss protonema cells (de Keijzer et al 2023).

In this study, we have generated transgenic *P. palmivora* lines expressing GFP-tagged α-tubulin and used these lines for live cell imaging of the microtubule cytoskeleton. We imaged dynamic microtubule processes including microtubule organization in the hyphae and microtubule behaviour during mitosis. We report microtubules to radiate astrally from MTOCs, with longer microtubules lying parallel to the hyphal axis. Microtubules radiating from MTOCs associated with adjacent nuclei established anti-parallel connections that maintained well beyond mitosis. Microtubules originating from the most apical MTOC reached into the hyphal tip. Depolymerization of microtubules caused erratic nuclear positioning, suggesting a mechanism of microtubule-mediated nuclear positioning in the coenocytic hypha, and disruptions in sustained hyphal growth.

## Results

### Expression of GFP-tagged α tubulin allows live cell visualization of microtubule organization and dynamics in *Phytophthora*

In the *P. infestans* genome, we identified five gene models encoding an α-tubulin subunit (van den Hoogen 2018). At the protein level, *Phytophthora* α-tubulins are highly similar (**figure S1**). For live-cell imaging of the microtubule cytoskeleton, we generated *P. palmivora* transformants expressing GFP-tagged α-tubulin, an approach that has been successfully implemented to visualize the microtubule cytoskeleton in a wide range of organisms (Goodson et al 2010). In the transformation constructs a GFP coding sequence followed by the open reading frame of either PITG_07960 (named *PiTubA2*) or PITG_07999 (*PiTubA5*) were inserted between the promoter and terminator the *Bremia lactucae* HAM34 gene (**figure S2**). Transformation of *P. palmivora* strain P6390 resulted in a total of six independent transformants with a detectable fluorescent signal. In all six lines - three for each of the two transformation constructs that were designated GFP-TubA2#1-3 and GFP-TubA5#1-3, GFP fluorescence showed a similar localization pattern reminiscent of microtubules (**figure S3**). We used the transgenic lines GFP-TubA2#1 and GFP-TubA5#1 for further experimentation. These lines behaved similarly as the wild type recipient strain in growth assays with no changes in viability and growth morphology (**figure S4**).

### Dynamic microtubules emanate from microtubule organizing centers

Previously it was shown that the microtubule cytoskeleton of *P. infestans* and *S. ferax* is organized in cytoplasmic microtubules (Temperli et al 1991) that originate from MTOCs (Heath et al 2000, Temperli et al 1990, Uchida et al 2005). These findings are in line with microtubule organization that we observed in this study in the *P. palmivora* GFP-TubA lines. Global analysis of microtubule localization revealed that hyphae possess MTOCs associated with the nuclear envelope that occur either individually or in pairs on each nucleus (**figure 1a**, (Evangelisti et al., 2019). In between paired MTOCs a spindle was often observed, indicating that the associated nucleus was undergoing a mitotic division (**figure 1a**). Short microtubules (1-2 μm) radiated into the cytoplasm from each MTOC in an aster-like organization, whereas axial microtubules (>2 μm) were oriented exclusively parallel to the long axis of the hyphae (**figure 1a**). Axial microtubules originating from MTOCs associated with adjacent nuclei appeared to interact now and then thereby forming connections between these nuclei (**figure 1a**). Axial microtubules originating from the most apical MTOC extended into the hyphal apex and appeared to polymerize against the apical cell membrane (**figure 1b**). The MTOCs from which they originated tracked the growing tip at fixed distances of 14.0 ± 2.1 µm (n = 43) (**figure 1b**), which is in the same range as the distance in between the most apical nucleus and the hyphal tip measured in *P. infestans* (Ketelaar et al 2012).

**Figure 1.**
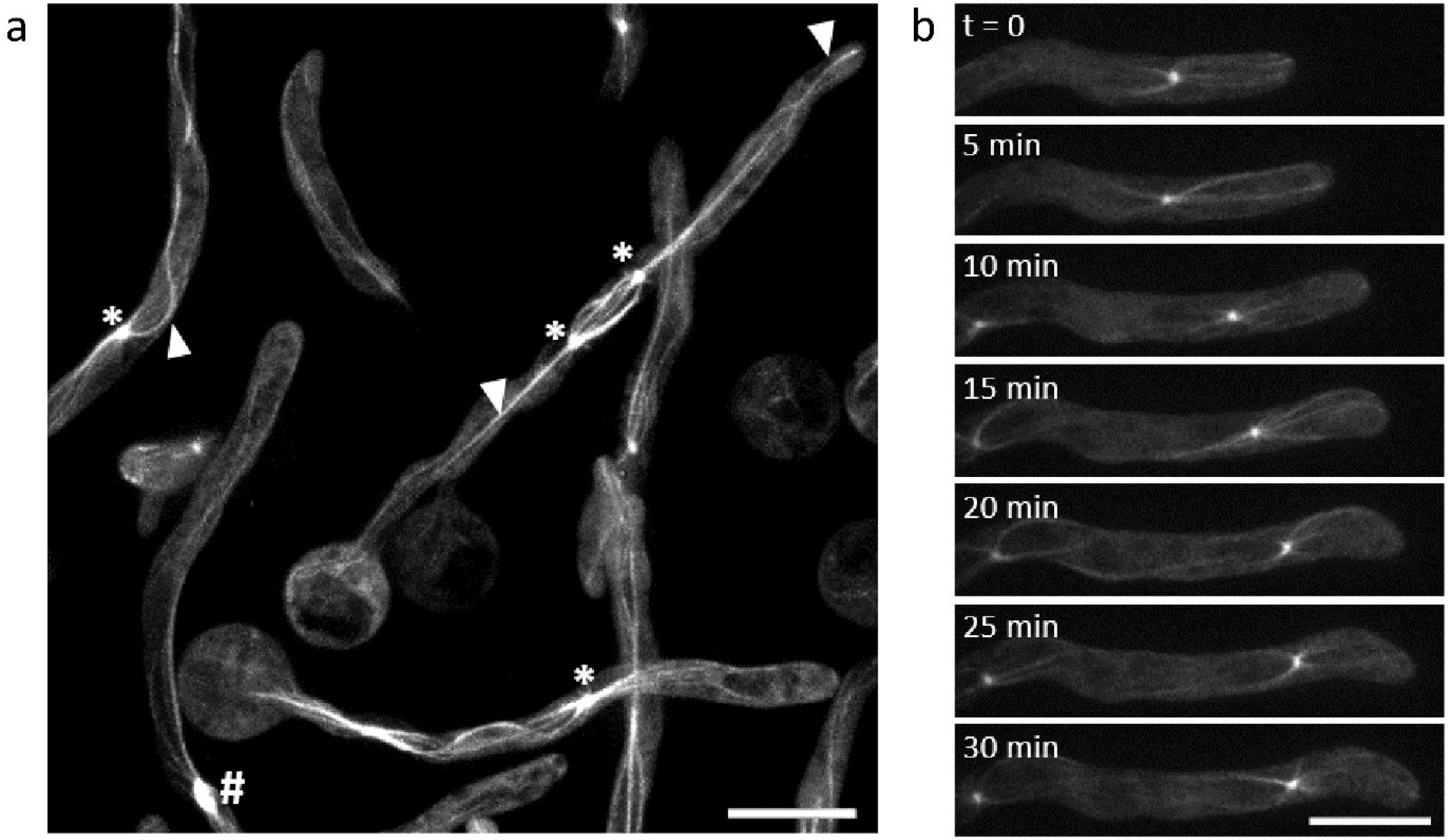
Microtubule organization in germ tubes of *P. palmivora* lines expressing GFP-tagged α-tubulin. **a**. Overview of microtubule organization in GFP-TubA2#1. Arrowheads indicate cytoplasmic microtubules. Asterisks highlight MTOCs and the hashtag indicates a spindle. **b**. Time series highlighting an example of dynamic microtubule reorganization in GFP-TubA5#1. Scale bars are 10 µm.

### Microtubule organization and dynamics in *P. palmivora* during mitosis

Oomycetes are coenocytes, having multiple nuclei that jointly reside in a shared pool of cytoplasm. Nuclei do not divide simultaneously and, as mentioned earlier, remain enclosed by the nuclear envelope during mitosis (Heath 1980). We observed microtubule dynamics during mitosis by monitoring germinating cysts of the *P. palmivora* GFP-TubA lines over time. During the initial stages of germ tube outgrowth, nuclear division occurs in a consistent and predictable manner, allowing us to track the microtubule organization during the progression of mitosis in *P. palmivora*. The first detectable sign of an imminent mitotic event was the duplication of the MTOC. After this duplication, both MTOCs were positioned at opposite sides of the nucleus. During this phase, that usually occurred within the first 30 minutes after zoospore encystment, spindle assembly did not occur yet (**figure 2a**). This stage lasted several hours.

**Figure 2.**
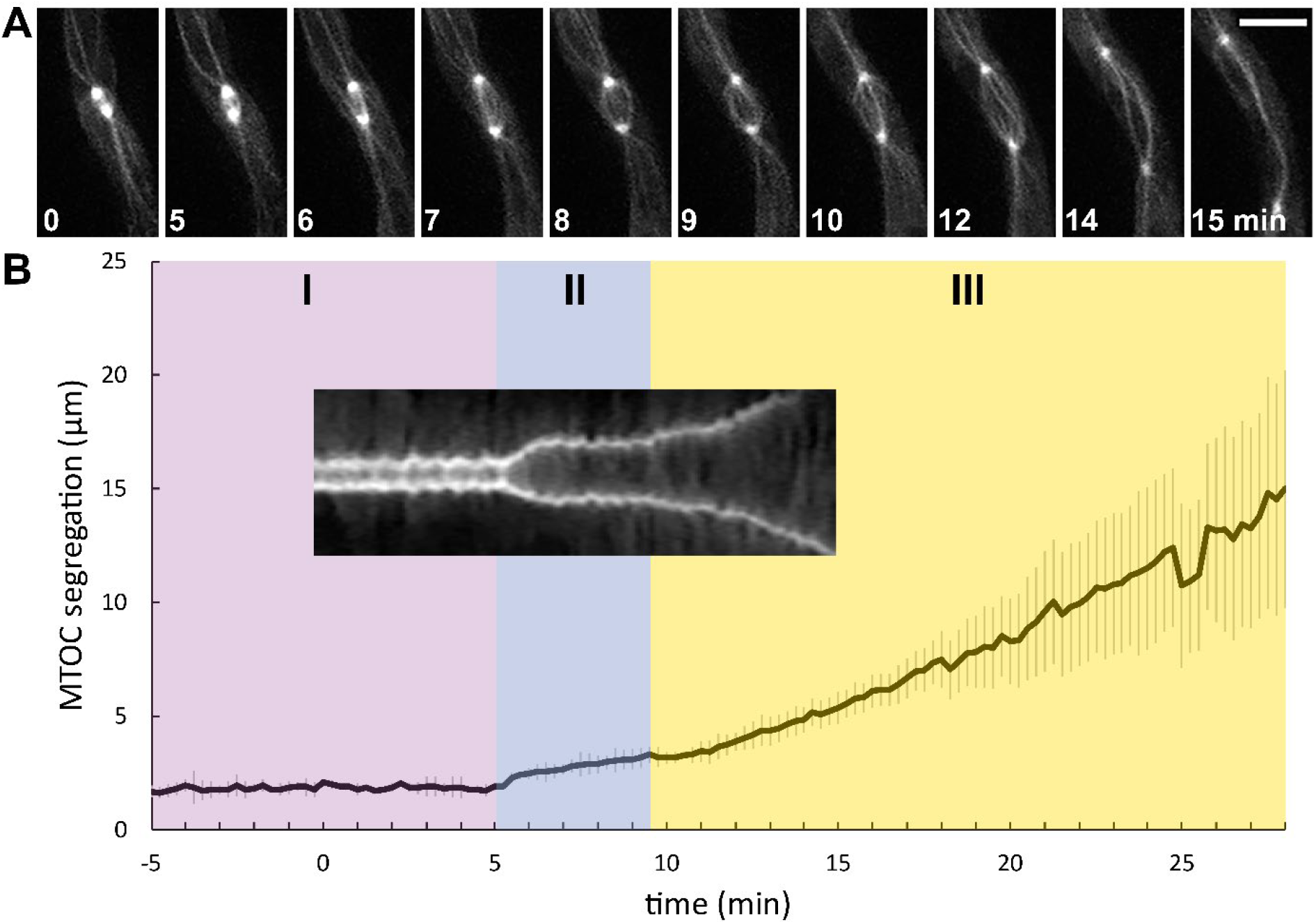
Microtubule organization during nuclear division in *P. palmivora*. Time series of **(a)** MTOC duplication and **(b)** mitotic spindle in hyphae of *P. palmivora* GFP-TubA5#1. c. Average distance (Y-axis) between the two spindle pole MTOCs during mitosis over time (X-axis) (n=11). Three different phases were identified based on the speed of microtubule segregation (I-III). t=0 was set at 3 hours after germination. Inset: Kymograph of spindle depicted in **(b)** showing the spacing between the MTOCs over time. Scale bars are 5 µm.

To track mitosis, imaging was initiated 3 hours after cyst germination with a focus on individual spindles to ensure sufficient spatiotemporal resolution. As the nuclear division progressed, distinct phases could be distinguished (**figure 2c**; I-III). During the first phase a microtubule spindle assembled while the MTOCs remained separated at a constant distance of 1.73 ± 0.19 µm (**figure 2c**; I). Overall, this stage persisted at least 30 minutes, and in some nuclei even over an hour. During the next phase (II), the distance between both MTOCs gradually increased with an average velocity of 13.45 (± 9.77) µm h^-1^. This stage lasted approximately 5 minutes. During the final phase (III), the separation of the MTOCs accelerated to a velocity of 41.95 (± 20.62) µm h^-1^ (**figure 2b**). Towards the end of this phase, MTOC segregation velocities decreased until a constant distance between the MTOCs was established (18.96 ± 6.15 µm, n = 5). To correlate MTOC dynamics to mitotic stages, we tracked nuclear dynamics during mitosis in a transgenic *P. palmivora* line named LILI-td-NT that expresses a nucleus-localized fluorescent protein mTFP1 (Evangelisti et al 2019), **figures S5 and S6**). This allowed us to correlate MTOC stage II to the anaphase, during which MTOC displacement is mediated by the elongation of interpolar microtubule pairs, referred to as anaphase B. During anaphase each pair of chromosomes is separated into two identical chromosomes. Stage III likely represents the telophase, during which MTOCs rapidly move apart and separation of the nuclear envelope occurs (**figures S5 and S6**). Nuclear separation -and consequently mitosis-is likely completed early on in stage III, as evidenced by the distinct and separated nuclear envelopes (**figures S5 and S6**).

### Cytoplasmic microtubules emerging from MTOCs interact with microtubules from adjacent nuclei and the hyphal tip

Anaphase B is a process during which, in vertebrate cells, the spindle pole MTOCs are separated by microtubule-based force generation; a combination of polymerization and microtubule sliding, during which overlapping microtubules are slid apart by motor proteins (Alberts et al 2023). Cytokinesis, which typically results in two daughter cells separating the divided nuclei and their associated microtubules, does not occur in the coenocytic hypha of oomycetes. Instead, microtubules originating from opposite MTOCs remain connected for an extended time frame after completion of mitosis. Since the - ends are associated with the MTOCs, the connection is likely the result of antiparallel interaction of the + ends. The intensity over the length of a putative microtubule pair is notably higher at the center, which is in line with the presence of an antiparallel overlap (**figure 3b**, blue arrowheads). Microtubules caught in these persisting connections were typically curved, indicative for a process referred to as ‘buckling’. Due to their rigid nature, buckling only occurs when microtubules are exposed to significant compressive forces (Kikumoto et al 2006). In older germ tubes with multiple nuclei, microtubules originating from adjacent MTOCs continued to remain connected beyond mitosis. Even in this stage microtubule buckling was frequently observed, often followed by subsequent displacement of the MTOCs (**figure 3a, b**). Besides these persistent interactions, we observed interactions between microtubules that were very short lived and resulted in rapid depolymerization of the interacting microtubules (**figure 3a**).

**Figure 3.**
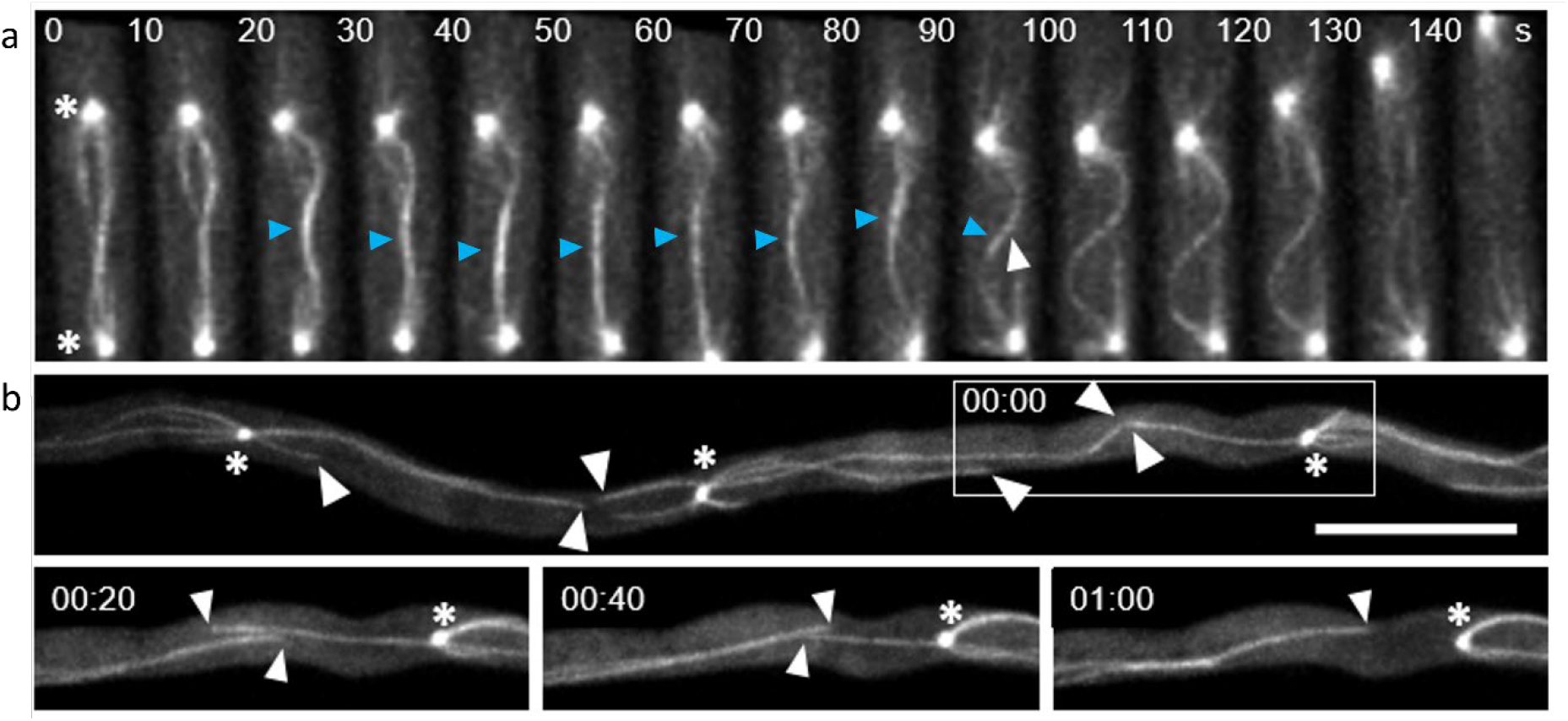
Dynamic behaviour of cytoplasmic microtubules in *P. palmivora*. **(a)** Polymerization and depolymerization of cytoplasmic microtubules in hyphae of *P. palmivora* GFP-TubA5#1. White arrowhead points to the dynamic plus-ends that alternate phases of polymerization and depolymerization, asterisks indicate two daughter MTOCs moving apart and blue arrowheads point to an area of increased intensity between two interacting microtubules, presumably forming an overlap. The width of the section at each time point is 5 µm. Note that the spacing between the MTOC increases during and after the buckling that occurs between 100-130 seconds. **(b)** Antiparallel connections between microtubules originating from adjacent MTOCs over time. Asterisks indicate MTOCs; white arrowheads point to buckling microtubules. Time is displayed in mm:ss; scale bar is 10 µm.

Besides microtubule-microtubule interactions, we frequently observed microtubule polymerization towards and close to the cell membrane (**figures 1 and 4a**). These interactions were most evident when axially oriented microtubules grew into the growing hyphal tip. Originating from the MTOC associated with the nucleus located adjacent to the hyphal tip, microtubules radiated into the apex and occasionally made contact with the apical cell boundary. Buckling events were specifically associated with microtubules that encountered the cell boundary at the hyphal tip (**figure 4a**; compare arrowheads 0 s and 120 s). To test if compressive forces are generated by the interaction of microtubules with the cell boundary, we assessed bending of microtubules that extended into the hyphal tip. This was done by calculating the ratio of the actual length of the microtubule extending into the apex to the shortest possible path length. In addition, we determined if the analysed microtubules interacted with the cell apex or not. This analysis showed that microtubules buckled significantly more (1.08 ± 0.08, n = 26) when they interacted with the cell cortex than those that did not (1.01 ± 0.08, n = 17; **figure 4b**). Moreover, the presence of a buckling microtubule at the tip was frequently followed by either displacement of the MTOC away from the tip or an arrest of its movement toward the tip (n=26; **figure 4c**). In contrast, displacement of the MTOC toward the tip was observed only infrequently (n=2).

**Figure 4.**
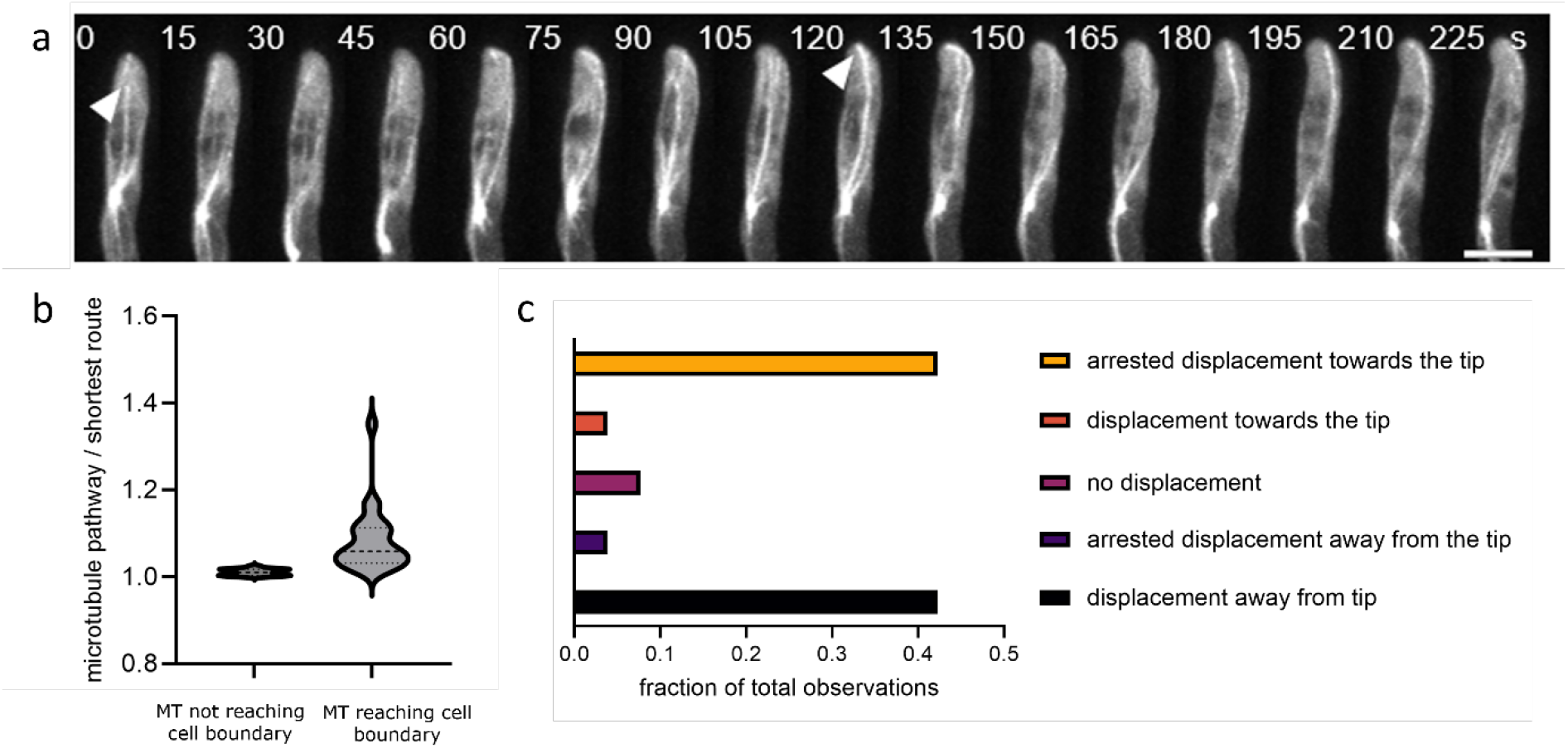
Cytoplasmic microtubules extend into the hyphal tip. **(a)** Microtubules extend into the apex of the growing hyphal tip of *P. palmivora* GFP-TubA5#1. In the 0s time frame, the arrowhead indicates the tip of a microtubule that is not in contact with the apical cell membrane. In the 120s time frame, the arrowhead indicates a microtubule that is in contact with the apical cell membrane. Scale bar is 5 µm. **(b)** Microtubule buckling increases in prominence during interaction of the tip with the apical cell membrane. The numbers in the graph represent the ratio of the microtubule length from the MTOC to its tip over the shortest path length from MTOC to tip; higher numbers indicate more buckling (P<0.05; two-sided t-test; n-26). **(c)** The presence of a microtubule that reaches the cell apex correlates with subsequent displacement of the MTOC away from the apex or arrest of apically directed motion (n=26).

### Microtubule depolymerization disrupts nuclear positioning

In oomycetes, MTOCs are associated with nuclei (Evangelisti et al 2019, Heath et al 2000, Temperli et al 1990, Uchida et al 2005) and presumably physically linked (Reinsch & Gönczy 1998). We observed a correlation between microtubules extending into the hyphal apex, microtubule buckling and subsequent distal MTOC displacement (**figure 4**), indicating that forces generated by these microtubules determine the position of the most apically located nucleus. This raises the question if the microtubules that connect MTOCs associated with adjacent nuclei play a role in the dynamic positioning of MTOCs and their associated nuclei. In several organisms nuclear positioning is known to be mediated by the microtubule cytoskeleton (Reinsch & Gönczy 1998), a feature that might be evolutionarily conserved in *Phytophthora*.

To investigate the putative role of the microtubule cytoskeleton in nuclear positioning in oomycetes, we exposed germ tubes of *P. palmivora* GFP-TubA lines to the microtubule depolymerizing drug oryzalin and found that oryzalin at concentration of 5-10 µM was sufficient to completely depolymerize cytoplasmic microtubules (**figure S7**). In contrast, MTOCs and spindles remained intact even at higher concentrations of oryzalin, suggesting that spindle MTs are rather resistant to depolymerization, possibly because the nuclear envelope somehow acts as a barrier for the drug (Heath et al 1984). Since cytoplasmic microtubules are the most likely candidates for nuclear positioning, we considered the effects of 5-10 µM oryzalin adequate to investigate microtubule-mediated nuclear positioning.

To investigate the effect of microtubule depolymerization on nuclear dynamics, we also exposed the *P. palmivora* LILI-td-NT line (Evangelisti et al 2013) to oryzalin (**figure 5**). In mock treated hyphae nuclei migrated along with the growing tip (**figure 5a**), with the distance between nuclei being relatively uniform (34.3 ± 10.9 μm, n = 1498, N = 29 hyphae, **figure 5b**). Hyphae treated with 10 µM oryzalin exhibited a significantly larger variation in nuclear spacing (36.3 ± 14.6 μm, n = 1118, N = 21 hyphae, **figure 5b, c**). Whereas in mock treated hyphae, nuclei maintained a fixed distance from the growing tip (23.8 ± 9.6 μm, n = 338, N = 29 hyphae), in oryzalin treated hyphae this distance was significantly increased and displayed a larger variation (39.0 ± 17.5 μm, n = 404, N = 21 hyphae, **figure 5d**).

**Figure 5.**
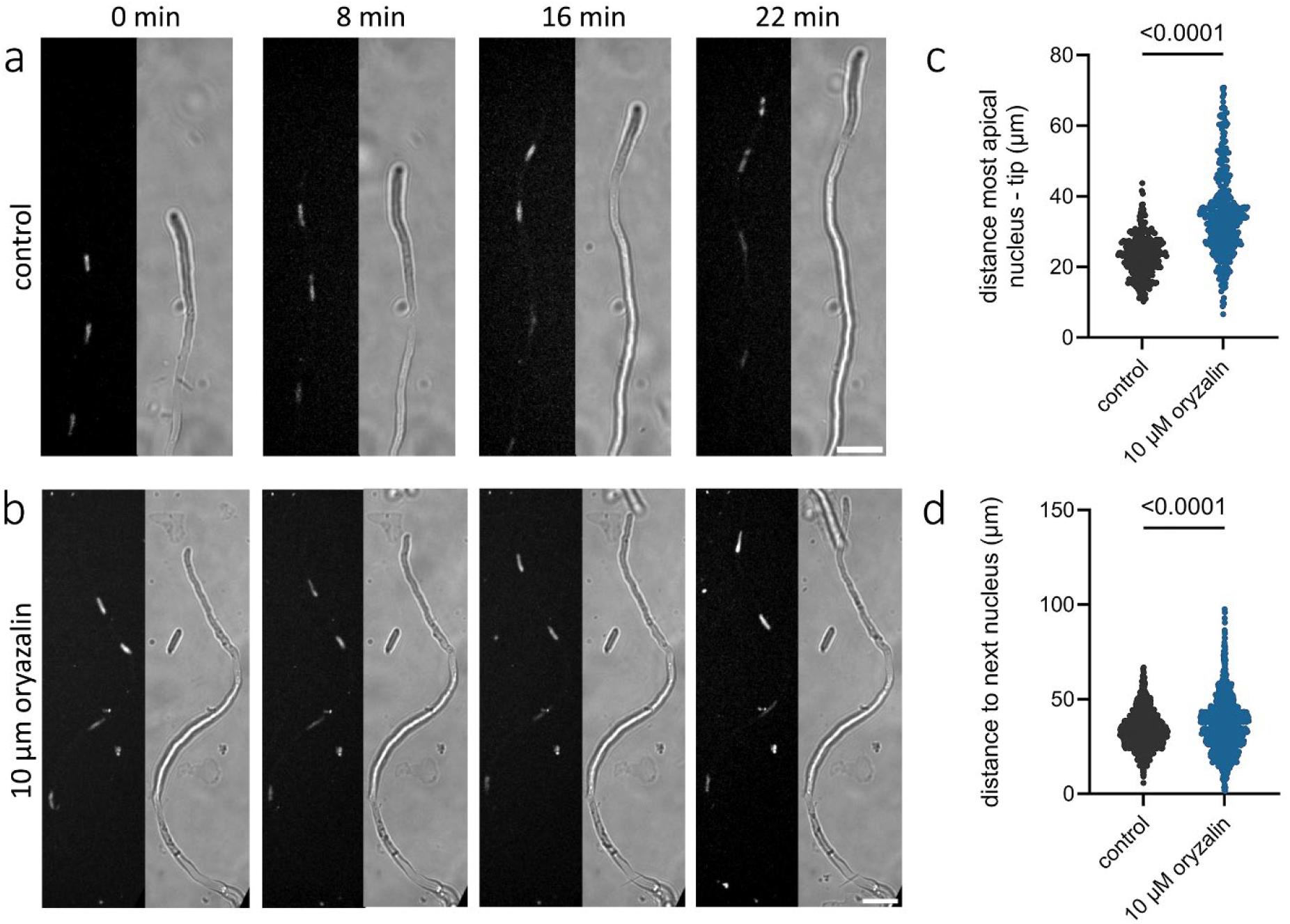
Effect of microtubule depolymerization on nuclear positioning in germ tubes of *P. palmivora* line LILI-td-NT. **(a,b)** Nuclear positioning over time during tip growth in (a) mock treated germ tubes and (b) germ tubes exposed to 10 µM oryzalin. Oryzalin was added 5 hours after the start of cyst germination. The right part of each panel shows the brightfield image. Debris in the medium originates from the V8 growth medium. Scale bars 20 µm. **(c**,**d)** Effect of 10 µM oryzalin on (**c**) internuclear distance (P<0.0001 by a two-sided t-test) and (**d**) distance between hyphal tip and the subapical nucleus (P<0.0001 by a two-sided t-test).

### Microtubule depolymerization disrupts sustained hyphal growth

When studying the role of microtubules in nuclear positioning, we occasionally observed hyphal growth defects. These hyphae we excluded from our analysis (**figure 5**) but prompted us to ask if microtubules function in hyphal growth. Microtubule depolymerization in tip growing cells typically leads to growth defects, which vary among different groups of organisms depending on the specific role of microtubules in tip growth. Functions include facilitating vesicle delivery to the growing tip, organizing the growth machinery, maintaining cell polarity, and orienting growth direction (Gudimchuk & McIntosh 2021, Horio 2007, Sieberer et al 2005). To investigate the role of microtubules in hyphal growth in *Phytophthora*, we studied the effect of microtubule depolymerization on germ tubes emerging from cysts (**figure 6**). In the presence of 10 µM oryzalin, the percentage of cysts that germinated as well as hyphal growth velocities were significantly reduced. Untreated hyphae elongated at an average velocity of 32.8 ± 16.7 µm h^-1^ (n=15) versus an average of 9.24 ± 8.06 µm (n = 15) in oryzalin-treated hyphae. Besides a reduced growth velocity in stretches of sustained growth, the oryzalin-treated hyphae also regularly showed growth arrests (0.76 ± 0.15 events h^-1^, n = 15), while all observed untreated hyphae showed sustained growth. In 28% of the cases, the growth arrest was temporal and hyphal growth resumed after a pause. In the other cases, growth arrest was permanent. We further noticed that the oryzalin-treated hyphae displayed excessive branching (0.64 events per tip h^-1^, n = 15, **figure S8**) whereas untreated hyphae did not branch at all in the 5-h timeframe during which we tracked growth. Branching appeared to be correlated with growth arrests: the frequencies of branching and growth arrests were similar (0.61 ± 0.22 events h^-1^, n = 15), and branching hardly occurred during phases of active hyphal growth (0.05 ± 0.09 events h^-1^, n = 15). These observations suggest that branching represents a repair mechanism that reinitiates growth and show that microtubules are important for sustained growth of *Phytophthora* hyphae.

**Figure 6.**
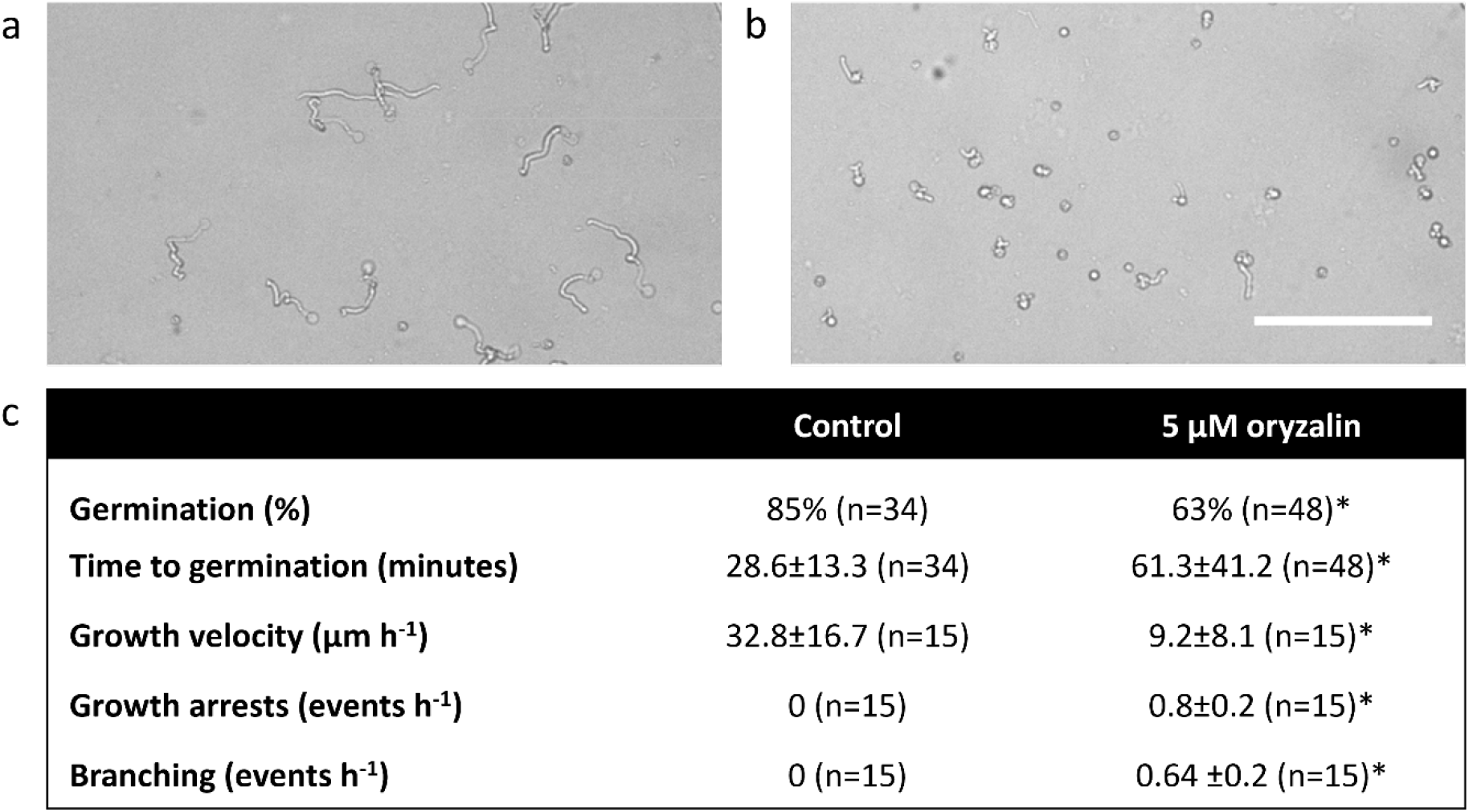
The effect of microtubule depolymerization on cyst germination and growth velocity of germ tubes. **a, b**. Progression of germination of *P. palmivora* GFP-TubA5#1 cysts after 5 hours in the absence (a) or presence of 5μM oryzalin (b). Scale bar indicates 15μm. c. Effects of oryzalin treatment on the listed parameters. Germination percentages were determined after 5 hours incubation; the other parameters were calculated over the 5h interval. Growth velocities were only calculated for uninterrupted stretches of hyphal growth. ^*^ indicates significance (P<0.05; two-sided t-test)

## Discussion

In this study we show that sustained antiparallel interactions between microtubules originating from MTOCs associated with adjacent nuclei function in maintaining internuclear spacing in coenocytic *Phytophthora* hypha. Microtubules from the most apical nucleus that reach into the hyphal apex, distance this nucleus from the cell tip. In addition, the microtubule cytoskeleton is essential for sustained tip growth. Microtubule organization through MTOCs associated with the nuclear membrane occurs in multiple eukaryotic kingdoms and is thought to be the ancestral machinery for microtubule organization (Yubuki & Leander 2013). This ancestral organizational control by a nucleus-bound MTOC (Dogterom & Yurke 1997) appears conserved in oomycetes (Heath & Greenwood 1968, Heath et al 2000, Temperli et al 1990, Uchida et al 2005). Our results show that the link between microtubules, MTOCs and nuclei are exploited for dynamic positioning of nuclei in the oomycete coenocytic body.

Microtubule polymerization generates powerful forces on a cellular scale in the piconewton range (Dogterom & Yurke 1997). In single-nucleate cells, the microtubule cytoskeleton has been shown to drive nuclear positioning by either polymerization of astral microtubules against obstacles or tethering to membrane-bound anchors (Attrill et al 2024, Hyman & White 1987, Takahashi et al 2001, Tissot et al 2017, Tran et al 2001). Oomycetes, however, are not single-nucleate. *Phytophthora* zoospores encyst, germinate and form germ tubes in which nuclei undergo mitosis in the absence of cytokinesis, resulting in a multinucleate organism known as a coenocyte. A coenocytic body plan supposedly allows for efficient nutrient exchange throughout the organism, and coenocytes can maintain genetic diversity within a single cell body (Marleau et al 2011, Sperschneider et al 2023). However, being a coenocyte puts demands on processes such as wounding responses and intracellular organization. A balanced intracellular organization includes the proper distribution of nuclei throughout the coenocytic body, which demands stringent control of nuclear positioning. Diverse organisms with multinucleate life stages employ their microtubule cytoskeleton to position nuclei in a shared cytoplasm. This is seen in syncytial muscle cells (Folker & Baylies 2013), in syncytia in the slime mould *Dictyostelium* (Tikhonenko et al 2016), in the *Drosophila* embryo (Tissot et al 2017), in coenocytic filaments in the yellow-green alga *Vaucheria* (Takahashi et al 2001) and in multinucleate fungal hyphae (Xiang & Plamann 2003).

We observed persisting interactions between microtubules from MTOCs associated with adjacent nuclei. These interactions either persisted from mitotic events or originated from new encounters during interphase (**figure 2**). Since the - ends of microtubules remain associated with MTOCs, the + ends interact in an antiparallel fashion. Studies in various organisms, including plants (Gaillard et al 2008; de Keijzer et al 2017), yeasts (Janson et al 2007, Maddox et al 2000, Straube et al 2003), mammals (Bieling et al 2010), fruit flies (Sharp et al 1999) and frogs (Nguyen et al 2018), have established diverse roles for antiparallel overlaps in microtubule organization and cellular functioning. In general, an antiparallel overlap is established when microtubules from opposite poles encounter.

MAP65/Ase1/PRC1 is a conserved class of proteins involved in establishing and maintaining such interactions (Kosetsu et al 2013; Kapitein et al 2008; Jagrić et al 2021). These proteins are recruited to antiparallel microtubule encounters, link both microtubules and serve as a scaffold to recruit secondary factors (Kosetsu et al 2013; Kapitein et al 2008; Jagrić et al 2021). Indicative of the presence of an antiparallel overlap between microtubules is an increased intensity of the GFP-tubulin at this location, which we also observed (**figure 3**).

Antiparallel bunding proteins recruit diverse proteins, including kinesin motor proteins to the overlap zone (refs). These proteins have functions in modulating polarization dynamics, reinforcing bundling and generating sliding forces, collectively dictating the dynamics of both microtubules caught in the antiparallel overlap (refs). In a coenocytic body, antiparallel interactions between microtubules originating from adjacent nuclei have been demonstrated to be part of the machinery capable of positioning nuclei: in a fission yeast mutant with multinucleate cells, positioning of nuclei was dependent on microtubule crosslinking proteins (Teapal et al 2021). We report that a similar positioning mechanism is employed to position nuclei in the *Phytophthora* coenocytic hypha. Although the involvement of MAP65/PRC1/Ase1 proteins in antiparallel microtubule overlap formation has not been studied in oomycetes, the key protein in establishing and maintaining these overlaps, MAP65/PRC1/Ase1, is conserved in the oomycete lineage (van den Hoogen 2018). Further research will have to decipher the precise mechanisms that establish these overlaps for controlling nuclear spacing. The frequent buckling events that we observed, make it likely that nuclear positioning is the result of locally generated forces that are integrated to position multiple nuclei.

We found microtubules to reach into the growing tip and observed that microtubule depolymerization leads to interrupted growth and branching. Do these microtubules in the tip have a function in increasing resilience of the growth machinery for sustained tip growth? A similar role for microtubules has been reported in tip growing cells of plants and fungi. In tip growing fission yeast cells, microtubules originating from MTOCs associated with the nuclear surface polymerize into the cell’s ends. Besides having a role in nuclear positioning, microtubules also deliver proteins to the cell cortex to establish polarity in the cell, such as the cell-end marker Tea1 in fission yeast (Sawin & Snaith 2004). In the filamentous fungus *Aspergillus nidulans*, the cell end marker TeaA is delivered to the hyphal tips by growing microtubules and is anchored there by TeaR. The cell-end markers position tip growth, and mutants lacking these cell end marker proteins display curved or zig-zag growing hyphae. Although this phenotype is distinct from the erratic growth upon microtubule depolymerization observed in this study in *Phytophthora*, it may be caused by similar microtubule-based mechanisms that deliver cell-end markers to for sustained hyphal growth.

During tip growth in plant root hairs a similar microtubule connection between cell apex and nucleus has been reported (Ketelaar et al., 2002; Sieberer et al., 2004; Brueggeman et al 2022). In these cells, the nucleus and other machinery involved in cell growth, collectively referred to as ‘tip-growth unit’, is associated with the growing tip (Emons and Ketelaar, 2009). When the position of the nucleus is experimentally mislocalized, tip growth is arrested (Ketelaar et al 2002). Tip growth requires continuous supply of cell wall and plasma membrane materials, and the misallocation of the nucleus likely disturbs this supply and/or the delivery of polarity markers. Since cytoskeleton inhibitors affect both growth and nuclear positioning in root hairs, untangling both processes is challenging. However, both the actin and microtubule cytoskeleton have been implicated in nuclear positioning (Ketelaar et al 2002; Brueggeman et al 2022).

Together, the results presented in this study showcase a microtubule-based mechanism for nuclear positioning: nuclei are spatially positioned by microtubules emanating from nucleus-associated MTOCs, which interact antiparallelly with microtubules emanating from other nucleus’ MTOCs. In addition to these inter-nuclear microtubule connections, we show that sustained tip growth is disrupted in the absence of microtubules, which may be caused by the misallocation of nuclei, the failure to orchestrate polarity in the cell or a combination of these processes. The mechanisms that facilitate these processes have not yet been studied in oomycetes. More in depth investigation of the microtubule cytoskeleton in *Phytophthora* and its associated proteins could be a stepping stone towards novel oomycete-specific targets for disease control.

## Materials & Methods

### Bioinformatics

The genome sequence of *P. infestans* stain T30-4 (Haas et al. 2009) and the transcriptome sequences of *P. palmivora* strain P16830 LILI (Evangelisti et al., 2017) were screened for genes or transcripts encoding α-tubulin by BLAST searches. Alignment programs in Geneious (https://www.geneious.com/features/prime) were used for multiple sequence alignments.

### Plasmid construction and transformation

To obtain N-terminally GFP-tagged constructs for expression in *P. palmivora*, α-tubulin genes *PiTubA2* (PITG_07960) and *PiTubA5* (PITG_07999) were amplified from *P. infestans* strain NL88069 genomic DNA using primers PITG_07960_NotI_F, PITG_07960_AscI_R, PITG_07999_NotI_F, and PITG_07999_AscI_R (table S1). The respective PCR amplicons were cloned in pGFP-N (Ah-Fong & Judelson 2011) using the restriction sites NotI and AscI, resulting in constructs pGFP-07960 and pGFP-07999 (**figure S2**). The construction of plasmid pTORKm34GWR for expression **of a nucleus-localized mTFP1 has been described previously (Evangelisti et al. 2019)**.

Stable *P. palmivora* GFP-TubA transformants with constructs pGFP-07960 and pGFP-07999 were generated by PEG/CaCl_2_-mediated protoplast transformation of strain P6390 and stable *P. palmivora* LILI-td-NT transformants with construct pTORKm34GWR by zoospore electroporation of strain P16830 (LILI). Transformation protocols are described in Supplementary method 1.

### Strains, culture conditions, life stages and imaging

*P. palmivora* strain P6390 (McHau & Coffey 1994) and strain LILI (P16830, (Torres et al 2010)) were routinely grown on 10% V8 medium (10% V8 juice, 1 g/l CaCO_3_, 1.5% technical agar) containing 20 μg/ml vancomycin, 100 μg/ml ampicillin and 50 μg/ml amphotericin B at 25 °C under continuous light. For selection of transgenic lines the medium was supplemented with 25 µg/ml geneticin for *P. palmivora* GFP-TubA2 and GFP-TubA5 and 100 µg/ml for *P. palmivora* LILI-td-NT. For *P. palmivora* 6390, GFP-TubA2 and GFP-TubA5 zoospore release, 4 to 6 day old plates were flooded with V8 broth (10 ml per plate) and incubated in the light for 5 to 20 minutes. For *P. palmivora* LILI-td-NT, plates were incubated at 4 °C for 30 minutes, after which they were flooded with MilliQ water and incubated at 25 °C for 20 minutes. Zoospores were encysted by vigorous shaking for 5 minutes. Zoospore concentrations were diluted with V8 broth to the desired concentration for imaging (typically 1^*^10^4^ zoospores/ml) and subsequently 100 µl cyst suspension was pipetted in 35 mm glass bottom dishes (MatTek, Ashland, USA). Depending on the aim of the experiment cysts were allowed to germinate for 0-24 hours at 25°C. For the oryzalin assay, cysts were supplemented with desired concentrations of oryzalin (100mM stock in DMSO). In these assays control samples were treated with equal amounts of DMSO, never exceeding 1% of the total volume. Individual hyphal apex positions were registered at each timeframe Δt. Each new position zt+1 is then subtracted from previous position zt to obtain a traversed distance. Averaging over the total number of timeframes yield average growth speed per hypha. Significant differences are determined with two-sided t-tests. Microtubule organization, mitosis and nuclear organization were observed using a Roper (Evry, France) Spinning Disc Confocal System on a Nikon Eclipse Ti microscope using a 100×, 60x and 40x, respectively, Plan apo oil immersion objective (NA 1.4) and a 491 nm laserline. Z-stacks were collected with 0.5 μm Z-intervals. Images were analysed using FIJI (https://imagej.net/Fiji).

## Acknowledgements

We thank the Wageningen Light Microscopy Centre (WU) for the use of their facilities. This research was funded by the research programme Graduate School Green Top Sectors (MK - project GSGT. GSGT.2018.024) and the ALW-JSTP program (JvdH - project 833.13.002) which are financed by the NWO Science domain (NWO-ENW) of the Dutch Research Council (NWO) and by the Food-for-Thought campaign from the Wageningen University Fund (KK).

## Supplementary information

### Supplemental method 1. Transformation protocols

*P. palmivora* transformants of strain P6390 were generated using a modified version of earlier described methods (van West et al 1999). Germinating sporangia (10^5^/ml) were incubated on a large petri dish (ø 15 cm) containing 25 ml 10% V8 broth for 18 h at 28 °C in the dark. Mycelia were washed in MQ to remove sporangia and incubated in 0.8 M mannitol for 10 min to induce plasmolysis, and subsequently protoplasted by incubation in protoplasting buffer [0.4 M mannitol, 20 mM KCl, 20 mM MES pH 5.7, 10 mM CaCl_2_, 5 mg/ml CELLULYSIN (Sigma-Aldrich) and 10 mg/ml Lysing Enzymes from *Trichoderma harzianum* (Sigma-Aldrich)] for 30-45 minutes at room temperature in the dark. After removing residual mycelial fragments by filtration (50 µm mesh), protoplasts were pelleted by centrifugation (4 minutes, 700 g). The protoplasts were resuspended in MT buffer (1 M mannitol, 10 mM Tris-HCl pH 7.5) and after a second centrifugation step in MTC buffer (MT + 25 mM CaCl_2_), then diluted with MTC buffer to 1·10^6^ - 5·10^6^ protoplasts per ml. 700 µl of the protoplast suspension was mixed with 30 µg circular plasmid DNA in 50 µl MQ. After incubation for 10 minutes at room temperature, 700 µl of freshly prepared PEG solution (50% PEG-3350, 10 mM Tris-HCl pH 7.5, 25 mM CaCl_2_, sterilized by filtration) was slowly added to the DNA-protoplast mixture. Protoplasts were regenerated overnight at 28°C in 25 ml rye sucrose medium (Caten & Jinks 1968) containing 1 M mannitol, without antibiotics. Regenerated protoplasts were pelleted by centrifugation (5 minutes, 1000 g), resuspended, and plated on selective plates containing 25 µg/ml geneticin. Plates were incubated at 28°C in the dark. Colonies appeared within 4 days.

To obtain a *P. palmivora* strain expressing a nucleus-localized mTFP1, *Phytophthora palmivora* P16830 (LILI) was transformed by electroporation as described in (Evangelisti et al 2019). pTORKm34GWR was extracted from *Escherichia coli* Top10 cells using the NucleoSpin Plasmid DNA Purification Kit (Macherey-Nagel) following the manufacturer’s instructions. For plasmid DNA preparation, DNA concentration was measured using a NanoDrop spectrophotometer (Thermo Fisher Scientific). The quality of the preparation was assessed by 260/280 and 260/230 absorbance ratios. Zoospores were harvested from one-week-old *P. palmivora* mycelium grown on V8-agar plates. After a 30-minute incubation at 4°C, sterile water was added to release zoospores. The zoospore suspension was mixed with modified Petri’s solution (final concentrations: 0.25 mM CaCl2, 1 mM MgSO4, 1 mM KH2PO4, 0.8 mM KCl) and 20 µg of plasmid DNA. The mixture was used for electroporation using a Gene Pulser Xcell Electroporation System (Bio-Rad) with exponential decay settings (500 V, 50 μF capacitance, and 800 Ω resistance). After electroporation, zoospores were incubated in liquid V8 medium at 25°C for 8 hours with gentle shaking. Transformants were then selected on V8-agar plates supplemented with 100 mg/L geneticin (G418).

**Table S1.**
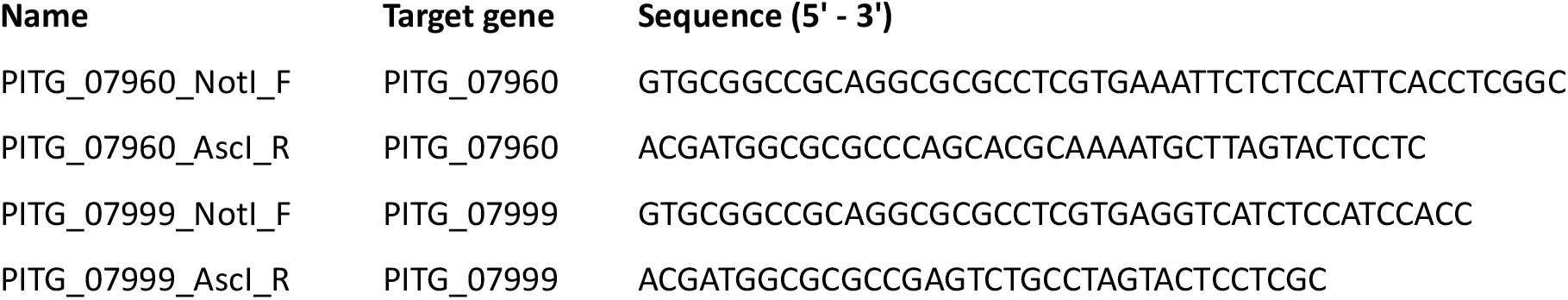
Primers used in this study.

**Figure S1.**
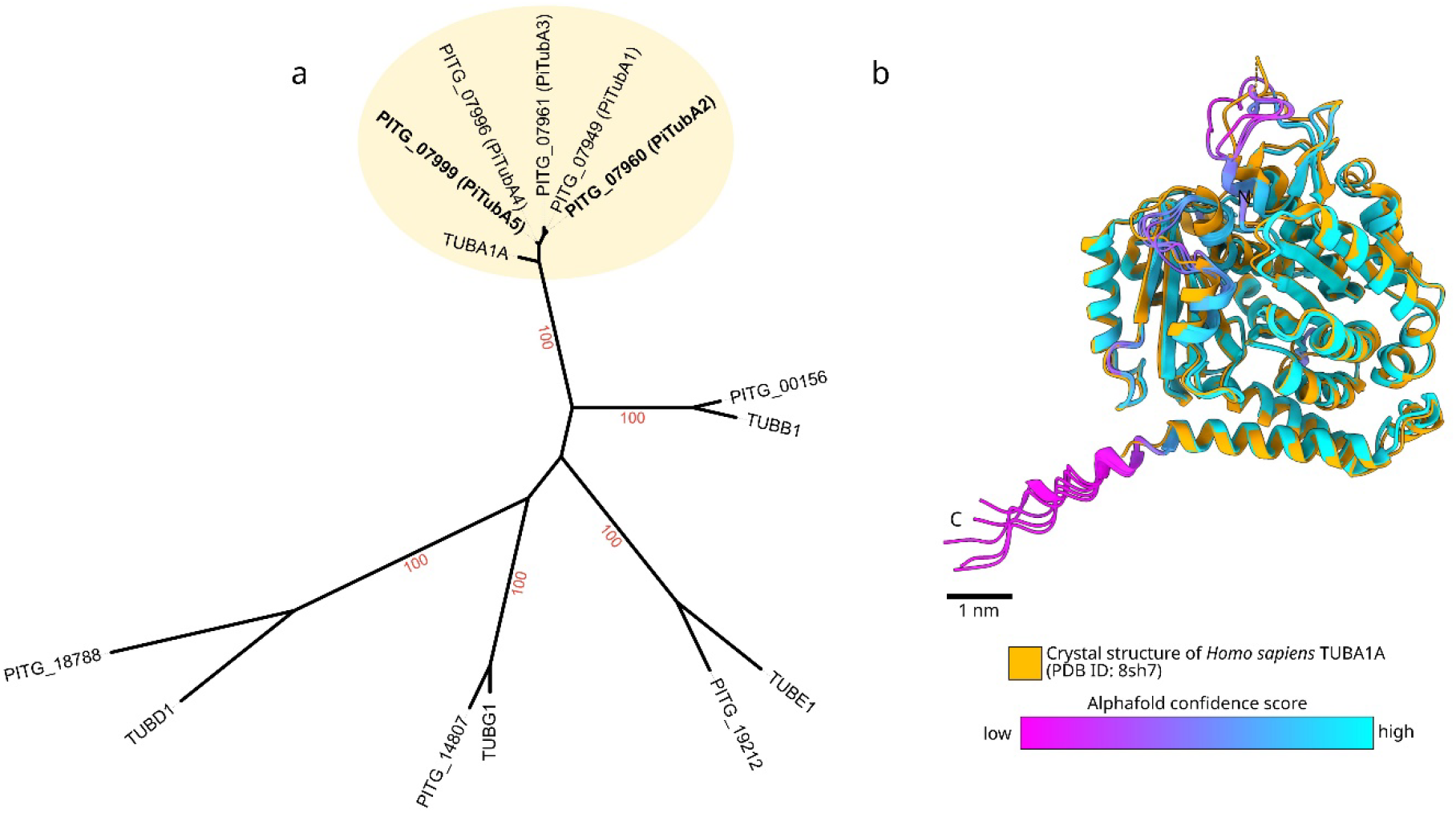
The *Phytophthora infestans* genome encodes multiple α-tubulin homologs. **(a)** Maximum likelihood phylogenetic tree showing the relationships between tubulin sequences from *P. infestans* (PITG gene models) and Homo sapiens. The clade containing *P. infestans* α-tubulins (PiTubA1– PiTubA5) and human TUBA1A is shaded in yellow. Bootstrap support values (in red) are shown at major nodes. **(b)** Structural comparison of *P. infestans* α-tubulin proteins predicted by AlphaFold (colored by confidence score: magenta = low, cyan = high) and the crystal structure of human TUBA1A (PDB ID: 8SH7, shown in orange). Structures are aligned to highlight conserved folding. Scale bar: 1 nm.

**Figure S2.**
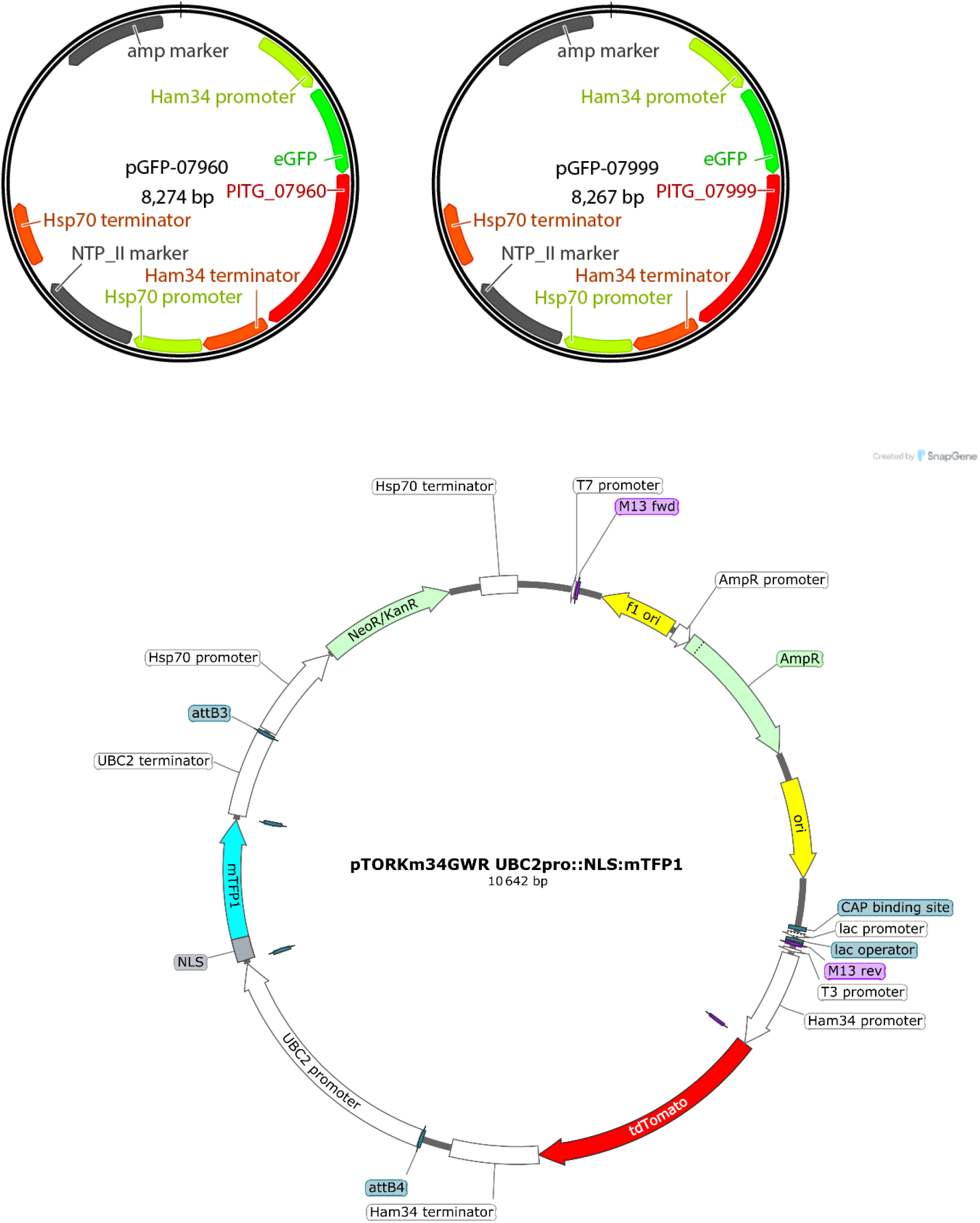
Transformation constructs. **a**. Constructs used for expressing N-terminally GFP-tagged *P. infestans* α-tubulins PITG_07960 (pGFP-07960) and PITG_07999 (pGFP-07999) in *P. palmivora* strain P6390. **b**. Construct used for dual labelling of nuclei and hyphae in *P. palmivora* strain P16830 (LILI).

**Figure S3.**
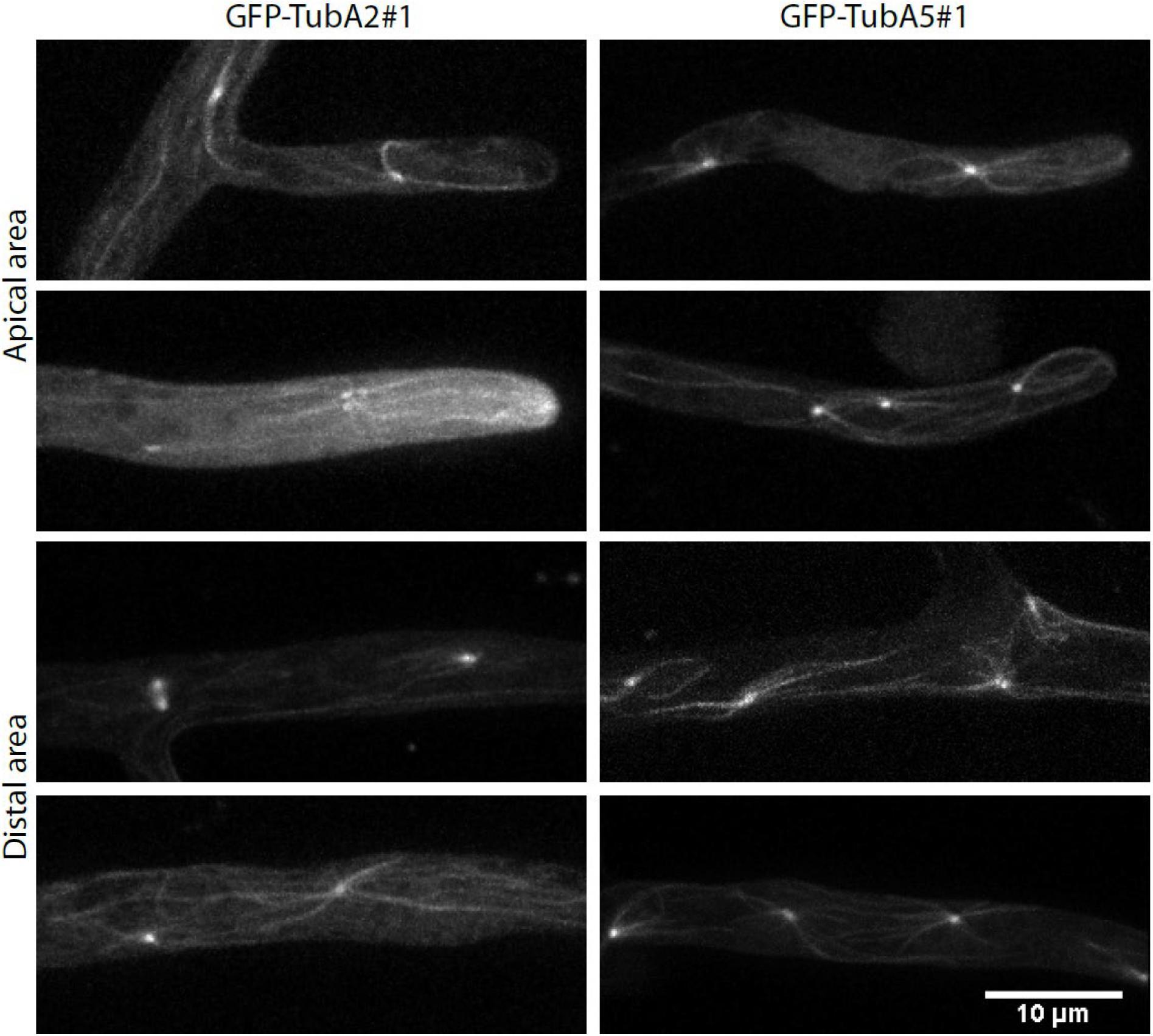
*P. palmivora* lines expressing GFP-TubA2#1 and GFP-PiTubA5#1 show similar localization patterns. In all lines, MTOCs, spindles and internuclear MTs were observed throughout the hyphae.

**Figure S4.**
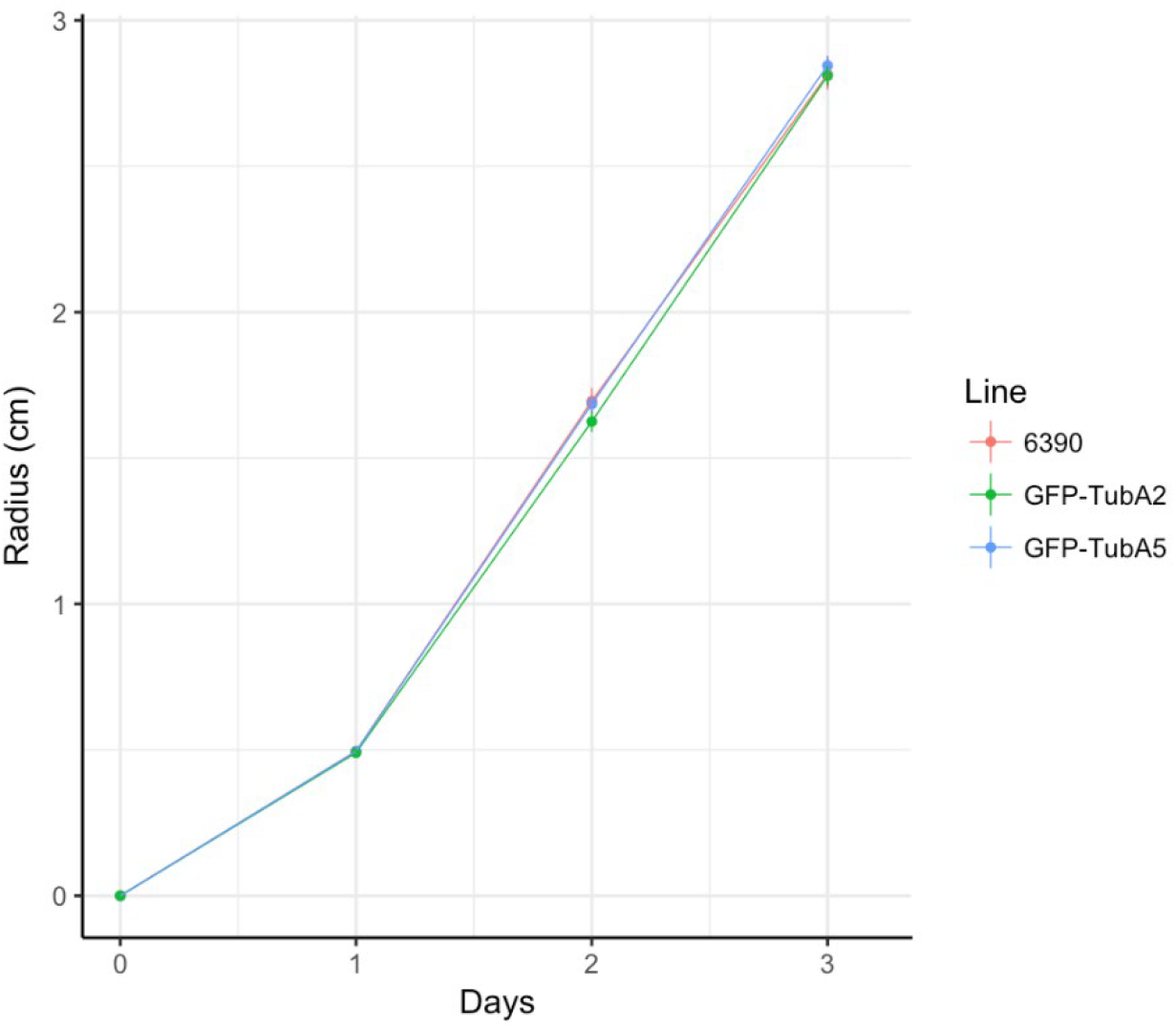
Radial colony growth rate of *P. palmivora* recipient strain P6390 (6390) and two GFP-tubulin strains (GFP-TubA2#1 and GFP-TubA5#1). The graph is representative for three independent growth rate assays that each included five replicates per line. Error bars represent standard deviations.

**Figure S5.**
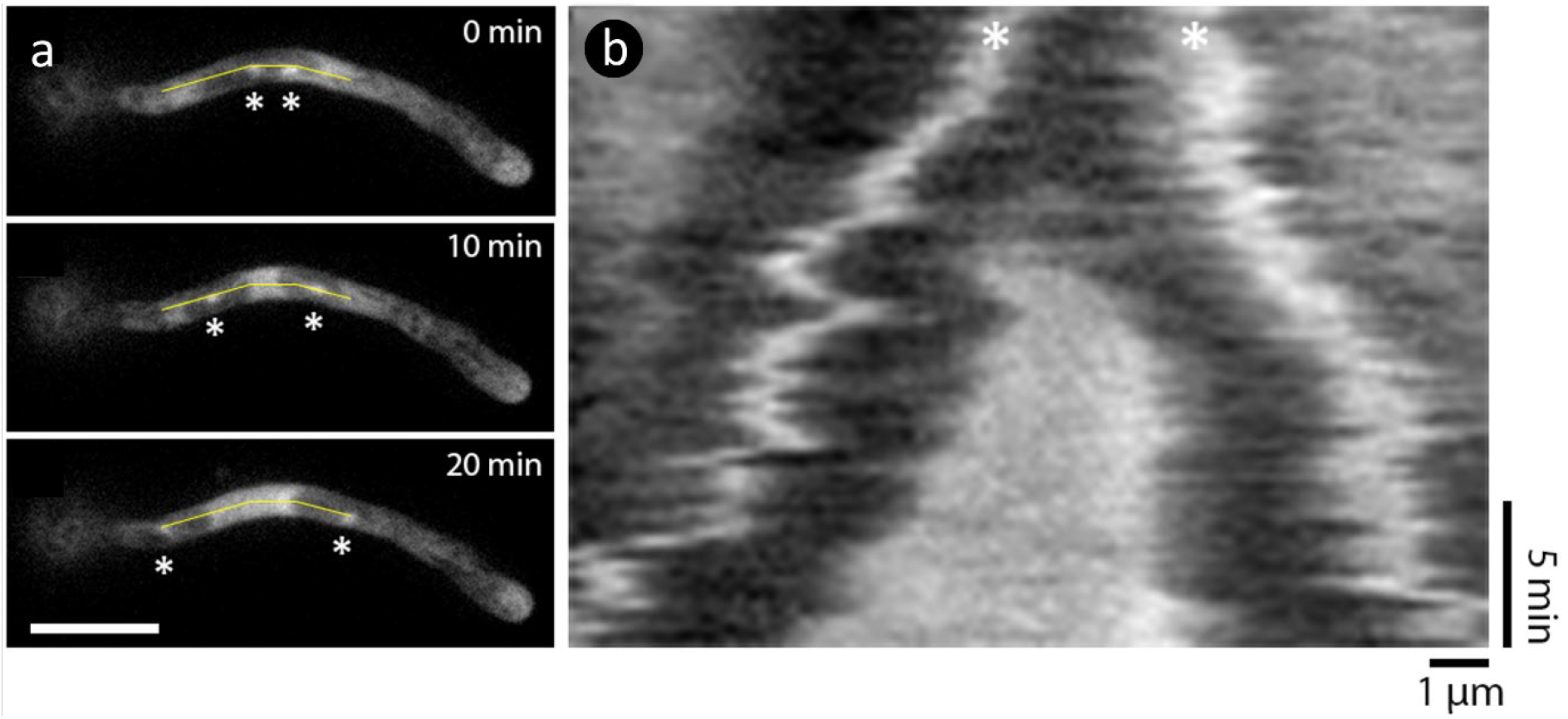
During mitosis, an area high in free tubulin appears between segregating MTOCs. **a**. Images of hyphae of *P. palmivora* GFP-TubA5#1. The line follows the path along which the MTOCs (indicated by ^*^) segregate during a 20 min time frame. **b**. Kymograph showing the movement of the MTOCs over time (20 min) and the appearance of a zone high in free tubulin.

**Figure S6.**
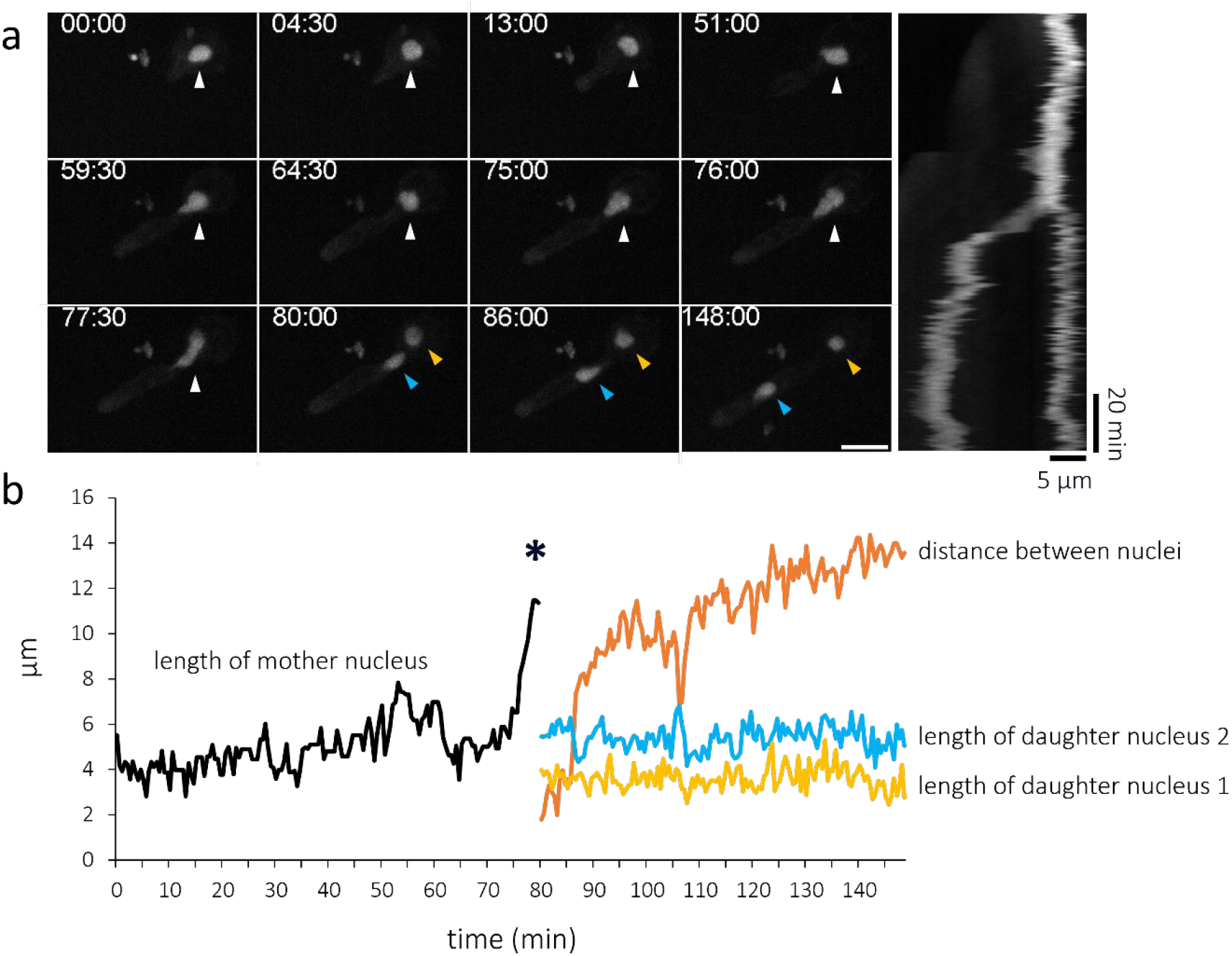
Mitosis in cyst and germ tube of a *P. palmivora* line expressing nucleus-localized mTFP (LILI-td-NT). **a**. Representative timeseries of a nucleus (white arrowhead) undergoing mitosis, resulting in two daughter nuclei (yellow and blue arrowheads). Scalebar 10 µm. Time in mm:ss. In the panel on the right: kymograph constructed along the long axis of the cyst-germ tube continuum. **b**. Length of nuclei and distance between separated nuclei (Y-axis) over time (X-axis). Graph shows mitosis progression of the nucleus shown in (a). Nuclear envelop separation (marked by ^*^) occurred at set length of the mother nucleus (11.37 ±2.54 µm, n=9).

**Figure S7.**
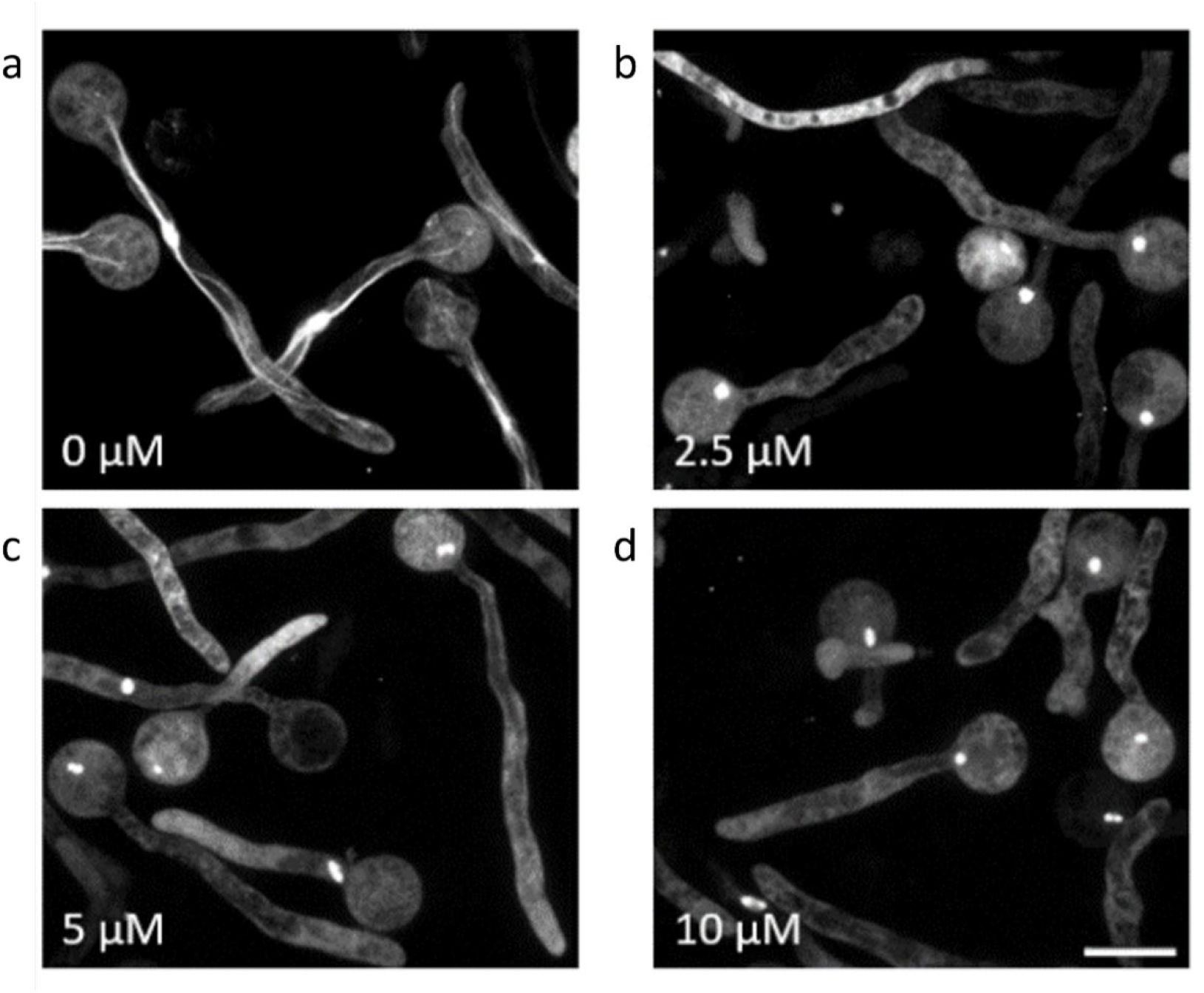
Oryzalin treatment depolymerizes cytoplasmic microtubules in germinating cysts of *P. palmivora* GFP-TubA2. Germinating cysts mock-treated (**a**) and treated with (**b**) 2.5 µM, (**c**) 5 µM and (**d**) 10 µM oryzalin. Scale bars: 10 µm.

**Figure S8.**
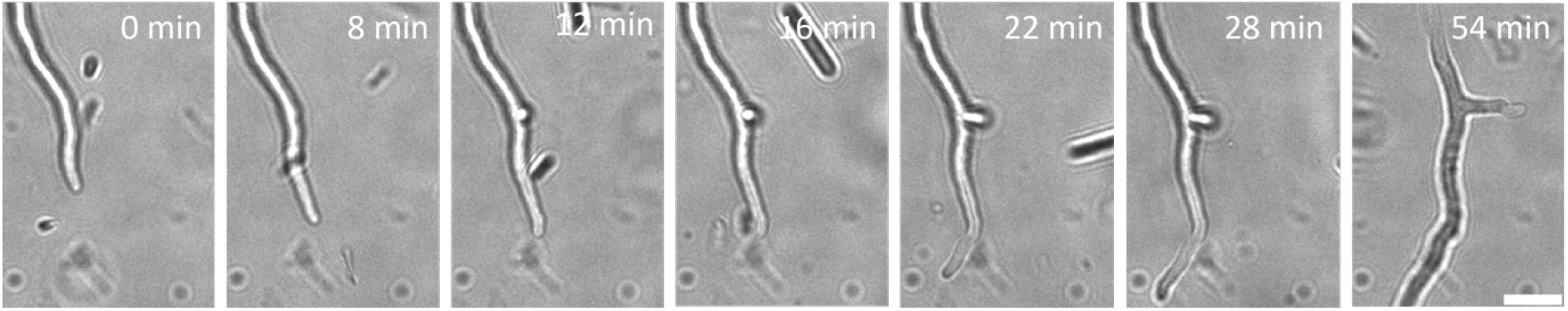
Growth arrest and subsequent branching event in a *P. palmivora* LILI-dt-NT hypha exposed to 10 µM oryzalin. Scale bar 15 µm.

